# Polarity signaling ensures epidermal homeostasis by coupling cellular mechanics and genomic integrity

**DOI:** 10.1101/401562

**Authors:** Martim Dias Gomes, Soriba Letzian, Michael Saynisch, Sandra Iden

## Abstract

Epithelial homeostasis requires balanced progenitor cell proliferation and differentiation, whereas disrupting this equilibrium fosters degeneration or cancer. Here we studied how cell polarity signaling orchestrates epidermal self-renewal and differentiation. Using genetic ablation, quantitative imaging, mechanochemical reconstitution and atomic force microscopy, we find that mammalian Par3 couples genome integrity and epidermal fate through shaping keratinocyte mechanics, rather than mitotic spindle orientation. Par3 inactivation impairs actomyosin contractility and viscoelasticity, and elicits mitotic failures that trigger aneuploidy, mitosis-dependent DNA damage responses, p53 stabilization and premature differentiation. Importantly, reconstituting myosin activity is sufficient to restore mitotic fidelity, genome integrity, and balanced differentiation and stratification. Collectively, this study deciphers a mechanical signaling network in which Par3 acts upstream of Rho/ROCK-mediated actomyosin contractility to promote intrinsic force generation, thereby maintaining mitotic accuracy and cellular fitness at the genomic level. Disturbing this network may compromise not only epidermal homeostasis but potentially also that of other self-renewing epithelia.

## INTRODUCTION

Epithelial sheets form important dynamic barriers between the interior of an organism and its environment, provide mechanical protection or mediate secretion, absorption and sensory functions. Being of central importance to survival, mechanisms have evolved to ensure that epithelial integrity is maintained during growth and quickly re-established upon injury. The epidermis is the major functional unit of the mammalian skin barrier and protects against harmful external insults or uncontrolled water loss^1^. In the developing epidermis, spindle orientation is coordinated with the specific need to produce either basal layer, self-renewing progeny or keratinocytes that commit to terminal differentiation and move suprabasally^2–4^. In adult epidermis, evidence emerges that also alternative mechanisms control cell fate decisions independent of spindle orientation. In ear skin, adjacent cells have recently been shown to undergo coupled, opposite fate decisions, with self-renewal being triggered by neighbor differentiation^5^. Moreover, accumulation of DNA damage has been implicated in HFSC loss and increased epidermal differentiation^6^, indicating that genome instability may serve as another parameter mediating tissue degeneration.

The function of many tissues also depends on a tight spatiotemporal control of cell and tissue architecture. Cell polarity refers to the unequal distribution of membrane compartments, organelles and/or macromolecules within a cell, and is critical for development, tissue homeostasis and repair^7,8^. Work in invertebrates identified the Par complex, consisting of *Par*titioning-defective3 (Par3), atypical protein kinase C (aPKC) and Par6, as a key module driving polarity processes including apico-basal polarity, asymmetric cell division and spindle orientation^4,9^. Recent work including ours showed that proteins of the Par3/aPKC complex control adult hair follicle stem cell (HFSC) maintenance and differentiation^10–12^. Altered spindle orientation was unlikely to cause these defects since Par3-deficient epidermis displayed a shift to more planar and oblique cell divisions^11^ instead of increased perpendicular divisions that are thought to fuel differentiation^2–4^. Therefore, we sought to investigate alternative molecular mechanisms through which polarity proteins mediate stem cell maintenance and control epidermal differentiation. Here, we provide evidence that Par3 controls epidermal integrity and fate predominantly through shaping Rho/ROCK-dependent keratinocyte mechanics, thereby sustaining the homeostasis of an important barrier-forming and self-renewing epithelium.

## RESULTS

### Epidermal Par3 loss leads to DNA damage responses

In search of potential mechanisms how epidermal deletion of *Par3* leads to progressive decline of HFSCs and ectopic epidermal differentiation^11^, we analyzed murine skin at postnatal day 100, a time-point right before Par3-deficient HFSCs start to decline^11^. Immunofluorescence analyses revealed increased signs of DNA damage throughout the skin epithelium of epidermal *Par3*KO (*Par3*eKO) mice, with a significantly increased number of γH2Ax-positive hair follicle bulges (Fig.1a,b, Suppl.Fig.1a), thus opening the possibility that DNA damage contributes to age-dependent loss of Par3-deficient HFSCs. In line with increased numbers of γH2Ax-positive cells, Par3 loss in keratinocytes resulted in a significant upregulation of the tumor suppressor p53 (Fig.1c,d, Suppl.Fig.1b), indicating genomic stress following *Par3* loss. We next assessed the activation of DNA damage checkpoint kinases earlier shown to activate and stabilize p53^13^. Intriguingly, loss of Par3-but not K14-promoter driven Cre expression alone-resulted in increased basal activity of the upstream checkpoint kinase ATR, and ectopic activation of both ATR and Chk1 kinases following irradiation with UV-B, a physiological stressor of epidermal cells (Fig.1e,f, Suppl.Fig.1c). Together, these data indicate that disturbed polarity protein function mounts ectopic DNA damage responses in HFSCs and epidermal keratinocytes.

### Par3 ensures mitotic fidelity and genome integrity

Searching for potential causes of elevated DNA damage signaling, we assessed the repair of damaged DNA lesions as Par3 has been implicated in repair of γ-irradiation-induced DNA double-strand breaks^14^. However, we did not obtain evidence for compromised DNA repair in Par3-deficient keratinocytes (Suppl.Fig.1d). Considering our earlier observations of altered spindle orientation in *Par3*eKO mice^11^ and the established link between aberrant chromosome segregation and DNA damage^15,16^ we next investigated the consequence of Par3 loss for cell division. Interestingly, time-lapse microscopy of H2B-GFP-expressing keratinocytes revealed a significant increase of aberrant cell divisions in Par3-deficient cultures that included lagging strands, multipolar divisions and cytokinesis failure (Fig.2a,b), accompanied by shortened mitotic duration (Fig.2c). Moreover, the mitotic defects correlated with a significant increase of aneuploidy in Par3-deficient k–eratinocytes as determined by interphase fluorescence in situ hybridization (iFISH) targeting chromosome 2 (Fig.
2d,e), providing a potential explanation for the observed p53 induction^17^.

**Figure 1.**
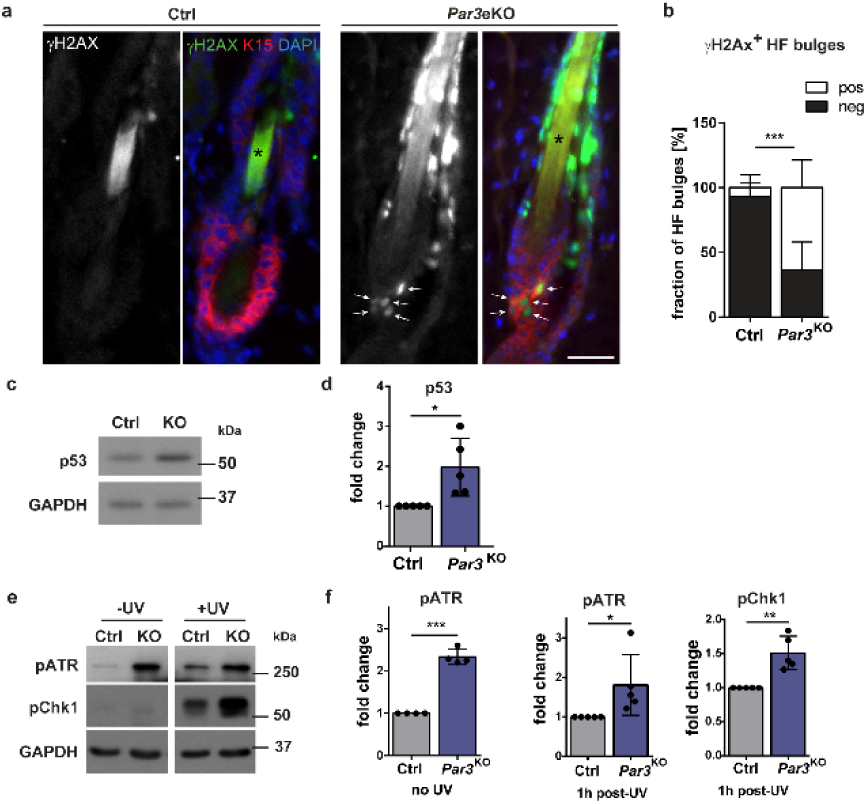
Loss of Par3 leads to increased DNA damage in hair follicle stem cells and ectopic activation of DNA damage responses. (a) Representative images of back skin sections from 100 days old epidermal Par3A knockout (*Par3*eKO: K14Cre/+;*Par3*fl/fl) and control mice (K14Cre/+) stained for γH2Ax indicative of DNA damage and Keratin15 to mark the bulge area. Scale bar: 25μm. Arrows mark γH2Ax and K15 double positive cells. Asterisks (*) mark unspecific staining of keratin in the hair. (b) Quantification of hair follicle bulges with cells positive for yH2Ax. n=5; mean±SD; two-way ANOVA/Sidak’s multiple comparisons test, ***: p<0.001. (c) Immunoblot for p53 total protein in whole cell lysates from primary keratinocytes isolated from *Par3*eKO and control mice. GAPDH was used as loading control. (d) Quantification of (c). p53 levels were first normalized to GAPDH (loading control) before values were normalized to control. n=5; two-tailed Student’s t-test, *: p=0.0171, mean±SD. (e) Immunoblot analysis of pATR and pChk1 in Par3KO and control keratinocytes either non-treated or UV-B treated (100mJ/cm2). GAPDH served as loading control. (f) Quantification of (e). pChk1 levels were first normalized to total Chk1, and pATR levels to GAPDH (loading control) before values were normalized to control. n(pATR, no UV)=4; n(pATR, pChk1 1h post UV)=5; two-tailed Student’s t-test, *: p=0.0457, **: p=0.0085, ***: p<0.001; mean±SD. Abbreviations used: HF, hair follicle; Ctrl, control.

Above findings unveiled an important role of Par3 for proper execution of mitosis and for maintaining genome integrity in mammalian epithelial cells. To directly assess if the mitotic defects were related to the elevated DNA damage responses observed in absence of Par3 we blocked mitosis using the Cdk1 inhibitor Purvalanol-A. Strikingly, Purvalanol-A treatment was able to significantly reduce the increased ATR activity in Par3-deficient cells (Fig.2f,g), supporting our hypothesis that erroneous mitosis following Par3 loss contributes to elevated DNA damage signals. Of note, compared to vehicle-treated cells, Purvalanol-A also led to a slight but non-significant reduction of ATR phosphorylation in control cells, potentially due to a baseline, Par3-independent mitotic inaccuracy (Fig.2b).

**Figure 2.**
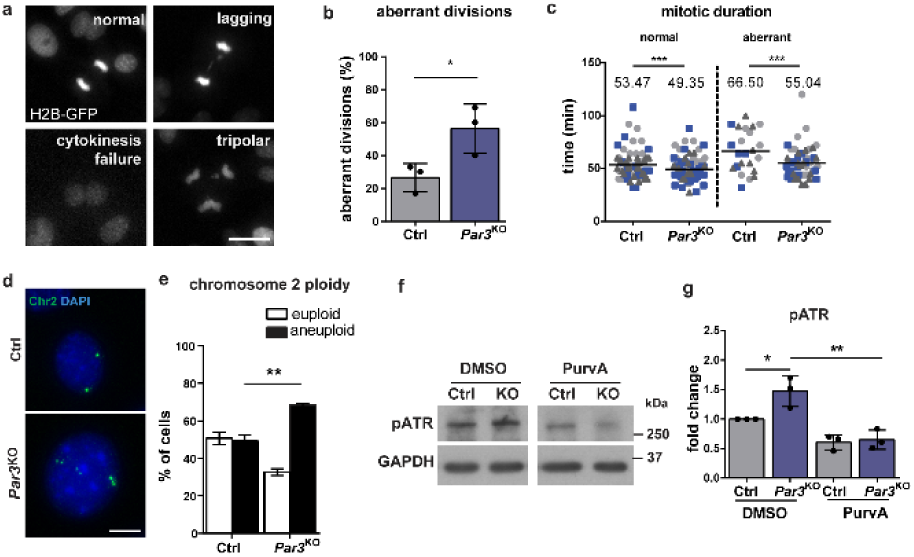
Par3 ensures mitotic fidelity and genome integrity of keratinocytes. (a) Representative images of mitotic aberration observed in time-lapse microscopy of H2B-GFP-expressing keratinocytes. Scale bar: 10μm. (b) Quantification of aberrant cell divisions in primary *Par3*KO and control keratinocytes observed in time-lapse microscopy. (c) Quantification of mitotic duration (Ctrl normal n=69, *Par3*KO normal n=95, Ctrl abnormal n=23, *Par3*KO abnormal n=51). For statistical analysis a nonlinear mixed model using R studio software was employed, yielding ***: p<0.001. Bar represents mean. (d) Representative images of iFISH probes targeting chromosome 2. Scale bar: 10μm. (e) Quantification of (d). For each experiment and condition at least 30 cells were counted. n=3; **: p=0.0028, mean±SD; two-way ANOVA/Sidak’s multiple comparisons test. Immunoblot analysis of ATR activation in primary Par3-deficient and control keratinocytes after treatment with Cdk1 inhibitor Purvalanol A. GAPDH served as loading control. (g) Quantification of (f). pATR levels were first normalized to GAPDH (loading control) before values were normalized to control. n=3; mean±SD; two-way ANOVA/Tukey’s multiple comparisons test *: p=0.0345; **: p=0.014. Abbreviations used: iFISH, interphase fluorescence in situ hybridization; Ctrl, control.

### Par3 safeguards contractility and keratinocyte dynamics

Having established that Par3 inactivation elicits disturbed mitosis and subsequent accumulation of DNA damage, we next aimed to understand the upstream events responsible for mitotic infidelity. First, the role of Par3 for overall cellular dynamics during cell division was examined employing detailed time-lapse microscopy of H2B-GFP-labelled keratinocytes followed by particle image velocimetry^18^. This analysis revealed a reduced strain rate of mitotic Par3-deficient keratinocytes as compared to control cells (Fig.3a,b), suggesting that Par3 orchestrates keratinocyte cell mechanics. During mitosis, cells undergo significant changes in shape, mechanics and polarity^19,20^ important for equal distribution of the genetic material^21,22^. Mitotic rounding, a ROCK-driven actomyosin-dependent process^22^, is important for bipolar spindle formation^23^. Par3 orthologs in C.*elegans* and Drosophila have previously been linked to actomyosin contractility^24–27^, albeit that the specific hierarchy among myosin activation and Par3 appears to be context-dependent. Based on the junctional localization of Par3 in keratinocytes^28^ we hypothesized that the mitotic aberrations could be caused by a primary defect in generating spatiotemporal contractile stresses at cell-cell contacts. Immunofluorescence analysis of primary keratinocytes indeed revealed a significant decrease of phosphorylated myosin light chain 2 (pMLC2) at intercellular adhesions (Fig.3c,d). In agreement, immunoblotting demonstrated a significant overall reduction in phosphorylation of MLC2 and of the actin-binding proteins ezrin–radixin–moesin (pERM) (Fig.3e,f), whereas total RhoA remained unaffected by Par3 inactivation (Suppl.Fig.2a,b). Consistent with this, immunohistochemistry analyses of murine skin demonstrated reduced phosphorylation of MLC2 in newborn and adult Par3-deficient epidermis (Fig.3g, Suppl.Fig.2c). Similarly, ERM phosphorylation, which in control mice was prominent in the epidermis and lower hair follicles, was reduced in adult *Par3*eKO mice (Fig.3h). These results suggested that Par3 is important to sustain mechanochemical signaling in epidermal keratinocytes *in vitro* and *in vivo*.

**Figure 3.**
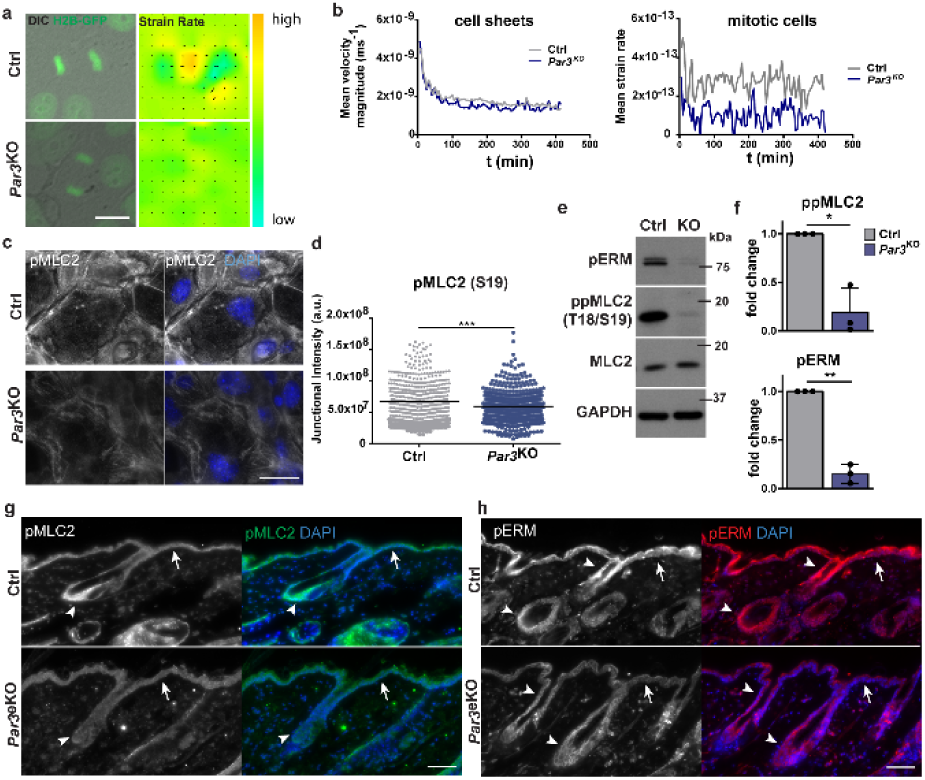
Par3 safeguards actomyosin contractility and keratinocyte dynamics. (a) Snapshots from live cell-imaging videos monitoring H2B-GFP fluorescence and DIC, and smoothened strain rate maps from particle image velocimetry analyses. Scale bar: 10μm. (b) Quantification of mean velocity magnitude (n=3 independent experiments) and mean strain rate over time (n=30 mitotic cells pooled from n=3 experiments). (c) Immunofluorescence micrographs of MLC2 phosphorylation (Ser19) in primary murine keratinocytes. Scale bar: 25μm. (d) Quantification of pMLC2 (Ser19) immunoreactivity at cell-cell junctions, intensity in arbitrary units; n = 450 cells pooled from three independent experiments; ***: p>0.0001, two-sided Mann–Whitney U test, bar represents mean. (e) Immunoblot analysis of ppMLC2 (Thr18/Ser19) and pERM. Total MLC2 and GAPDH were used as loading control, respectively. (f) Quantification of (e), ppMLC2 (Thr18/Ser19) levels were first normalized to MLC2 and then to GAPDH (loading control) before all values were normalized to control. n=3; two-tailed Student’s t-test, *: p=0.0299, mean±SD. pERM levels were first normalized to GAPDH (loading control) before all values were normalized to control. n=3, two-tailed Student’s t-test, **: p=0.004, mean±SD. (g,h) Immunohistochemistry of P100 murine back-skin paraffin sections stained for pMLC2 (Ser19) (g) and pERM (h). Scale bar: 75μm. Arrows point to pMLC2 and pERM signals in the interfollicular epidermis, arrow heads mark the upper and lower compartments of the hair follicle. Abbreviation used: Ctrl, control.

### Restoring myosin activity rescues force generation and elasticity upon Par3 loss

To examine the significance of reduced myosin phosphorylation for the mechanical defects observed upon Par3 loss, we next sought to restore myosin activity using a low dose of Calyculin-A, a potent phosphatase inhibitor and enhancer of actomyosin contractility. Protein biochemical analyses demonstrated that 1nM Calyculin-A indeed was able to rescue phosphorylation of MLC2 and ERM in Par3-deficient keratinocytes (Fig.4a). To functionally assess the role of Par3 in intrinsic force generation, epithelial sheet contraction assays were performed. After Dispase-mediated lift, Par3-deficient epithelial sheets showed significantly impaired contraction when compared to control cell sheets (Fig.4b,c), indicative of reduced internal stress. Strikingly, contraction of *Par3*KO keratinocyte sheets could be restored by treatment with Calyculin-A (Fig.4b,c). Similarly, direct activation of Rho using a small peptide stabilizing endogenous Rho-GTP was able to rescue contraction of *Par3*KO keratinocytes (Fig.4d), indicating that Par3 acts upstream of Rho/ROCK/myosin to mediate myosin motor activity and intrinsic contractile forces.

**Figure 4.**
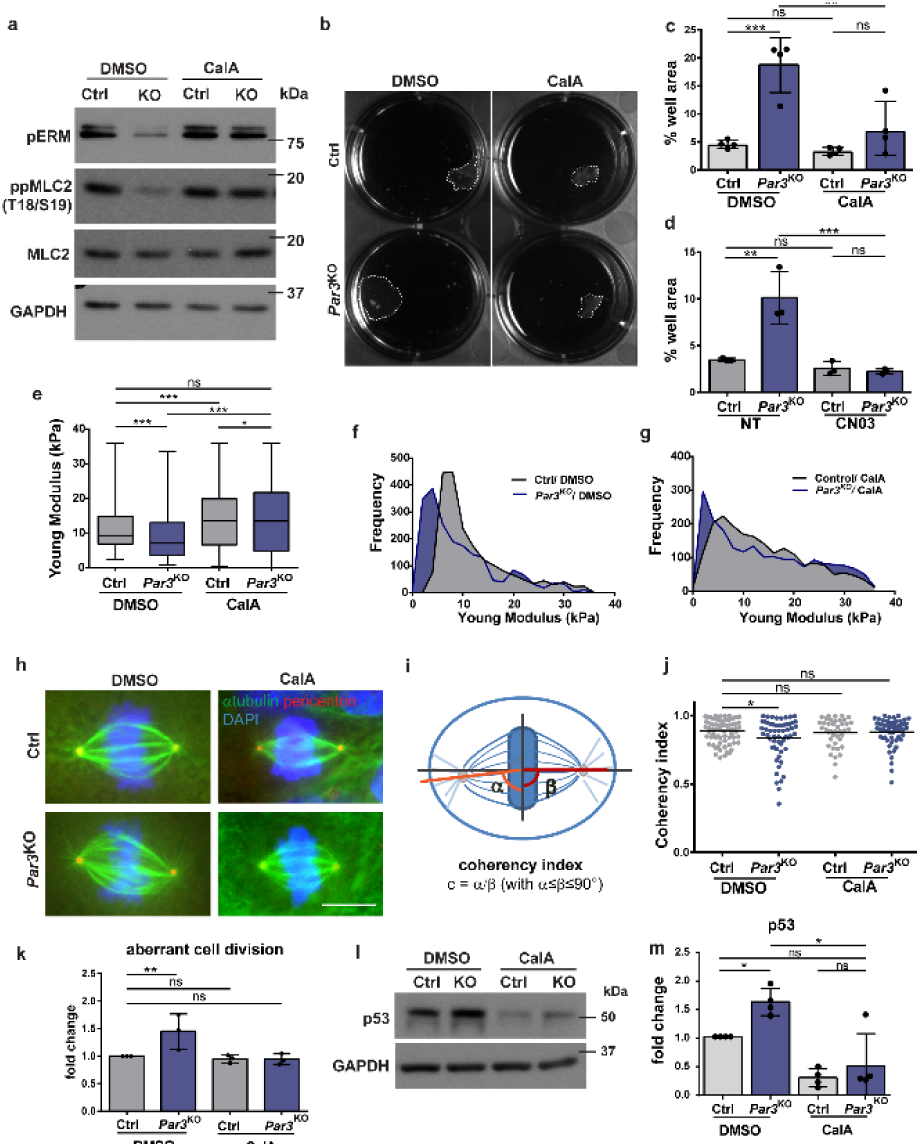
Re-establishment of actomyosin contractility rescues viscoelastic properties, mitotic fidelity and p53 upregulation in absence of Par3. (a) Immunoblot analysis of pERM and ppMLC2 (Thr18/Ser19). MLC2 and GAPDH were used as loading controls. (b) Representative images of dispase sheet contraction assay 24hours after 1nM CalA treatment using control and *Par3*KO keratinocytes. (c) Quantification of keratinocyte sheet area after dispase sheet contraction assay 24 hours following treatment with CalA. n=4; **, p=0.0029, ***, p = 0.0004, mean±SD; one-way ANOVA/Tukey’s multiple comparisons test. (d) Quantification of keratinocyte sheet area after dispase sheet contraction assay 24 hours following treatment with the Rho activator CN03 (5ng/ml). n=3; **, p=0.0024, **, p = 0.0011, ***, p=0.0008, one-way ANOVA/Tukey’s multiple comparisons test. (e) Young Modulus box-plot based on force indentation spectroscopy (n=2000 measurements, pooled from three independent experiments; *, p=0,038, ***, p=0.0002, Kruskal–Wallis/Dunn’s multiple comparison test, box plots show minimum, 25% percentile, median, 75% percentile and maximum). (f,g) Distribution histogram of Young Modulus upon DMSO (f) and 24h CalA treatment of experiments shown in (e). (h) Immunofluorescence micrographs of mitotic keratinocytes stained for pericentrin and alpha-tubulin, scale bar: 10μm. Intensity was enhanced for better spindle visualization. (i) Schematic representation of analysis for mitotic spindle geometry. (j) Quantification of coherency index (n=4 independent experiments; Ctrl DMSO treated: n=74, *Par3*KO DMSO treated: n=58; Ctrl CalA-treated: n=41, *Par3*KO CalA-treated: n=55), *, p=0.0458, mean, one-way ANOVA/Tukey’s multiple comparisons test. (k) Quantification of aberrant cell divisions in primary *Par3*KO and control keratinocytes observed in time-lapse microscopy upon DMSO and CalA treatment, n=3, ** p=0.0038, two-way ANOVA/Dunnet’s comparison test. (l) Immunoblot analysis of p53 upon DMSO and CalA treatment. GAPDH was used as loading control. (m) Quantification of (l). p53 levels were first normalized to GAPDH (loading control) before all values were normalized to controls. n=4; *: p=0.0440 (DMSO treated Ctrl vs. *Par3*^KO^); *: p=0.0318 (*Par3*KO DMSO-treated vs. CalA-treated); mean±SD; one-way ANOVA/Tukey’s multiple comparisons test. Abbreviations used: Ctrl, control; ns, non-significant; CalA, Calyculin-A.

As the actomyosin network is a major determinant of mechanical properties of cells and their responses to stress, we next employed atomic force spectroscopy to quantitatively assess the consequence of Par3 loss for keratinocyte mechanics. Interestingly, nano-indentation experiments using either sharp or spherical cantilevers uncovered impaired viscoelastic properties upon Par3 inactivation, with a significantly reduced elastic modulus measured in Par3-deficient keratinocytes compared to controls (Fig.4e,f, Suppl.Fig.3a). Similar to epithelial sheet contraction, Calyculin-A treatment of *Par3*KO keratinocytes was sufficient to rescue the elastic modulus (Fig.4e-g, Suppl.Fig.3b,c), thus demonstrating that Par3 regulates viscoelastic properties through control of actomyosin-dependent contractile behavior in epidermal keratinocytes.

### Contractility-dependent mitotic infidelity and p53 response

To elucidate how impaired actomyosin contractility contributes to mitotic fidelity, we first characterized mitotic spindles in terms of their intrinsic geometry. Intriguingly, whereas in metaphase control keratinocytes the two poles of a spindle exhibited the expected perpendicular orientation toward its metaphase plate, Par3-deficient cells frequently showed a deviation of spindle angles relative to the equatorial chromosome plate, reflected by a significantly reduced coherency index in these cells (Fig.4h-j). This was further associated with an increased spindle circularity in *Par3*KO keratinocytes (Suppl.Fig.3d). Strikingly, both types of metaphase spindle shape alterations could be normalized by Calyculin-A treatment (Fig.4j, Suppl.Fig.3d). Time lapse microscopy of H2B-GFP-labelled keratinocytes confirmed this was relevant for the outcome of mitosis as reconstitution of myosin activity in Par3-deficient cells was sufficient to re-establish mitotic fidelity (Fig.4k). In accordance with the observed mitosis-dependent DNA damage responses (Fig.2g), Calyculin-A treatment also significantly rescued the p53 upregulation in Par3-deficient keratinocytes to the levels of control cells (Fig.4l,m). Together, these data support a model in which insufficient contractility as consequence of Par3 inactivation leads to erroneous cell divisions and subsequent mounting of ectopic DNA damage responses.

### Low contractility upon Par3 loss drives ectopic differentiation and suprabasal fate

Evidence emerges connecting DNA damage and aneuploidy with reduced self-renewal capacity and increased differentiation in flies^29^ and mammals^30,31^. Recent studies in mouse skin and cultured keratinocytes associated DNA damage accumulation with keratinocyte differentiation^6,32^. Having demonstrated that myosin activation as well as inhibition of mitosis prevents ectopic DNA damage responses in absence of Par3, we hypothesized that the reduced genome integrity due to insufficient force generation fuels differentiation upon Par3 loss. To directly test this, we assessed the impact of restored contractility on expression of the epidermal differentiation markers Keratin1 and Involucrin following Ca^2+^-induced keratinocyte differentiation. Consistent with our previous findings of increased epidermal differentiation in *Par3*eKO mice^11^, immunoblot analyses confirmed significant Keratin1 and Involucrin upregulation in primary *Par3*KO keratinocytes compared to controls (Fig.5a,b, Suppl.Fig.4a). Strikingly, Calyculin-A treatment was sufficient to reverse this effect (Fig.5a,b, Suppl.Fig.4a), highlighting that impaired actomyosin contractility contributes to premature differentiation in the absence of Par3. Notably, similar results were obtained using ATM and ATR inhibitors (Suppl.Fig.4b), supporting our hypothesis that elevated DNA damage responses owing to impaired mechanical force generation contribute to increased differentiated fate upon Par3 ablation.

**Figure 5.**
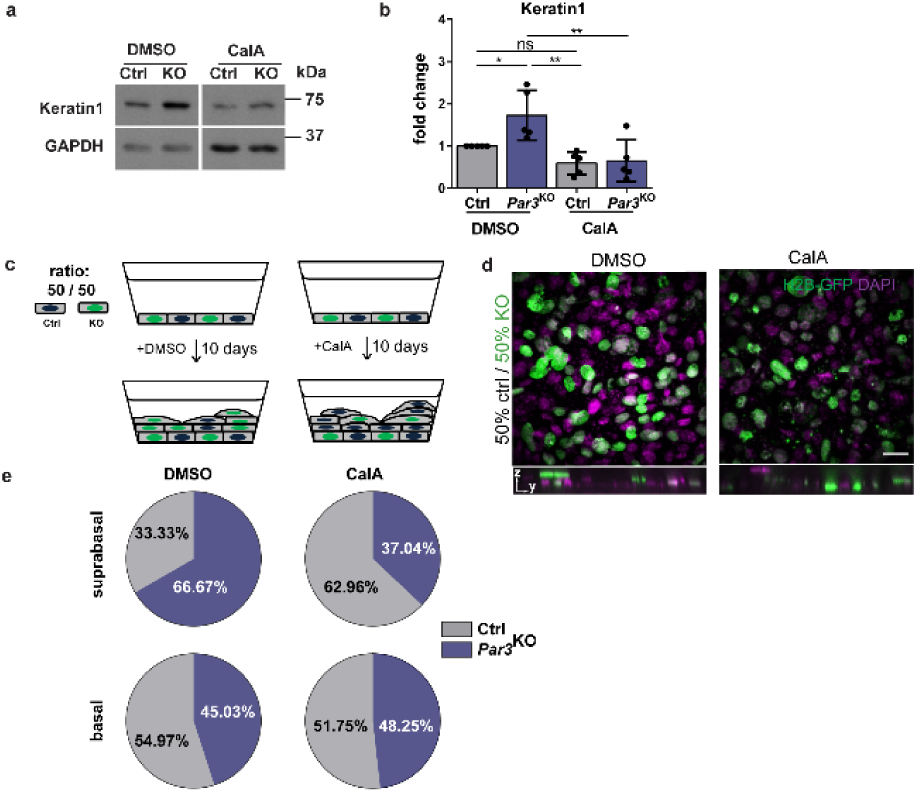
Reduced actomyosin contractility following Par3 loss drives ectopic differentiation and suprabasal fate. (a) Immunoblot analysis for Keratin1 expression in keratinocyte lysates following 48h CalA treatment. GAPDH served as loading control. (b) Quantification of (a). K1 levels were first normalized to GAPDH (loading control) before all values were normalized to controls. n=5; *: p=0.0431 (Ctrl vs. *Par3*^KO^, DMSO-treated); **: p=0.0034 (*Par3*^KO^ DMSO-treated vs. CalA-treated); **: p=0.0024 (*Par3*^KO^ DMSO-treated vs. Ctrl-CalA treated); mean±SD; two-way ANOVA/Tukey’s multiple comparisons test. (c) Schematic representation of the Transwell filter culture system and experimental setup. *Par3*KO keratinocytes were positive for H2B-GFP, control keratinocytes isolated from littermates were negative for H2B-GFP. (d) Immunofluorescence micrographs of stratified cultures incubated for 10 days either in DMSO (vehicle control) or CalA. Scale bar: 50μm. (e) Pie graphs showing cell distribution according to basal or suprabasal position following a 10-days culture period of mixed control and *Par3*KO keratinocyte cultures on permeable filters. Abbreviations used: Ctrl, control; ns, non-significant; CalA, calyculin.

To further corroborate our findings, we tested if the tight control of the actomyosin network was also crucial to direct keratinocyte fate toward upward movement into suprabasal layers. For this, primary keratinocytes were cultured for prolonged periods on porous membranes to enhance keratinocyte stratification into basal and suprabasal keratinocyte layers, better mimicking the *in vivo* tissue architecture. Primary control and *Par3*KO keratinocytes were mixed and seeded at a 50/50% ratio, using H2B-GFP as genetic label distinguishing genotypes (Fig.5c). After 10 days of culture stratification was achieved, and the number of control vs. *Par3*KO cells in basal and suprabasal layers was examined. These experiments revealed a consistent enrichment of Par3-deficient cells in the suprabasal layer relative to control cells (Fig.5c-e), regardless of the initial seeding ratio (data not shown), thus illustrating that *Par3*KO keratinocytes were more likely to move upwards, in line with a predisposition to differentiation. Most importantly, restoring actomyosin contractility by low dose Calyculin-A could revert the ratio of suprabasal control vs. *Par3*KO cells (Fig.5c-e), emphasizing that low myosin activity in Par3-deficient cells is a crucial driver of suprabasal, differentiated fate.

## DISCUSSION

In self-renewing tissues, stem/progenitor cell divisions need to be tightly controlled to maintain a genetically and quantitatively stable stem/progenitor cell pool while producing sufficient numbers of daughter cells that commit to differentiation and serve overall tissue function. Our biomechanical, genetic and imaging approaches uncover mechanisms through which the polarity protein Par3 instructs keratinocyte mechanics, mitosis and thereby ultimately epidermal fate decisions. We provide *in vivo* and *in vitro* evidence that Par3 acts upstream of Rho/ROCK/actomyosin activation to control epidermal homeostasis, illuminating a previously unrecognized hierarchy of polarity and mechanical signaling in the skin. Based on our findings we propose a model (Fig.6a) in which junctional Par3 safeguards genome integrity through control of actomyosin contractility and keratinocyte mechanics to ensure mitotic fidelity. Par3 inactivation leads to impaired myosin activity and causes mitotic infidelity, aneuploidy and DNA damage responses yielding p53 stabilization. Mechanistically, we demonstrate that mitosis is required to mount the increased DNA damage responses in Par3-deficient keratinocytes, and that restoring myosin activity is sufficient to normalize mitosis and to prevent ectopic DNA damage, premature differentiation and suprabasal fate.

**Figure 6.**
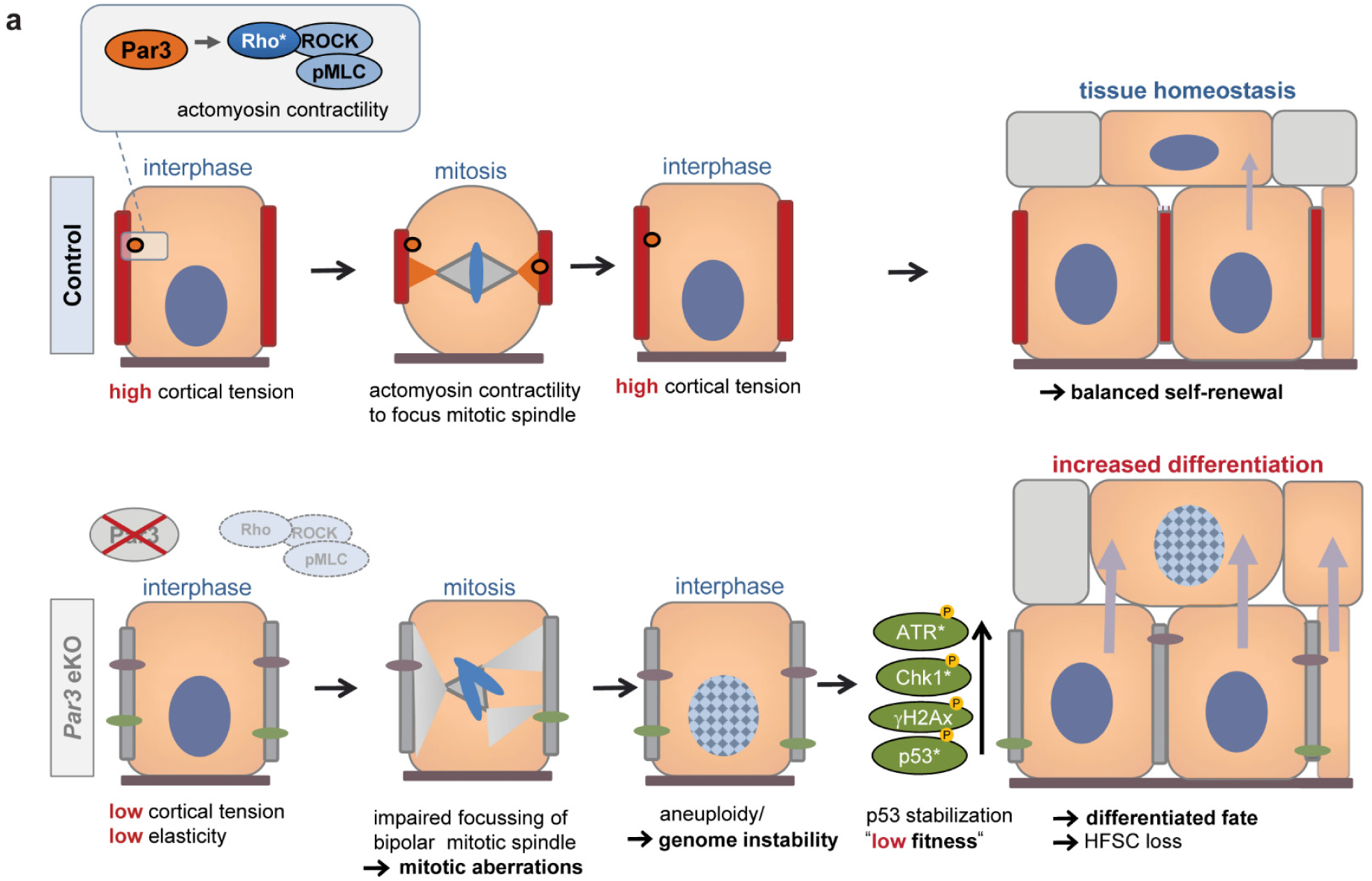
Par3 directs epidermal fate decisions through coupling actomyosin contractility to mitotic fidelity. (a) *Model*. Par3 regulates actomyosin contractility in a Rho/ROCK dependent manner. Loss of Par3 alters keratinocyte mechanics resulting in mitotic infidelity and accumulation of DNA damage and p53, which in turn fuels differentiation and disturbs epithelial homeostasis. This is associated with upregulation of P-cadherin and DDR1, previously implicated in suppression of actomyosin contractility. Restoring myosin activity in Par3-deficient keratinocytes is sufficient to normalize mitosis, and to prevent ectopic DNA damage, premature differentiation and suprabasal fate. This data establishes a novel role of Par3 in keratinocyte mechanics. Rather than promoting asymmetric/perpendicular mitotic spindle orientation, altered polarity protein function in adult epithelia causes ectopic differentiated fate due to impaired cortical contractility that yields mitosis-related DNA damage and as consequence lower “fitness”.

Our data further strongly suggest that genome instability and the resulting DNA damage responses act as defining factors for differentiated epidermal cell fate. These data are in agreement with recent developmental studies linking p53 to embryonic stem cell differentiation^33^ and data from mouse embryos exposing p53 as driver of differentiated fate by reducing cell competition and fitness^34,35^. Our findings extend these developmental data to a more general concept that orchestrates tissue homeostasis. In light of the frequent extrinsic and intrinsic damage that cells are facing over time, tissues may utilize biochemical alert signals reflecting compromised genome quality as a parameter for fate decisions to sustain long-term stem/progenitor cells with highest genome integrity, whereas cells exhibiting (moderate) DNA damage are eliminated by commitment to differentiation.

Contrasting the established role of Par3 in spindle orientation in several developmental systems^4,11,36,37^, our data indicate that polarity proteins help maintain adult tissue homeostasis primarily by controlling keratinocyte mechanics and thereby mitotic fidelity and genome integrity, rather than by sole orientation of mitotic spindles. Considering our new findings, it is tempting to speculate that, next to causing mitotic errors and DNA-damage triggered differentiation, disturbed cortical contractility also elicits the observed spindle orientation defects, e.g. due to improper spindle anchorage at the cortex, without that the latter necessarily impinges on cell fate. Such view of cortical tension acting upstream of LGN-NuMA localization and/or spindle orientation would be consistent with earlier observations made in epidermal Serum Response Factor-deficient mice^38^ and fly imaginal disc epithelial cells with suppressed actomyosin function^39^.

Par3 inactivation stabilizes surface^40^ and junctional P-cadherin (Suppl.Fig.2d,e), thus inversely correlating with actomyosin contractility. Despite the emerging role of cadherins in mechanotransduction^41^, the relevance of P-cadherin in mechanical force control is less well understood and may depend on its expression relative to E-cadherin^42,43^. P-cadherin can control mechanical forces at intercellular adhesions and during collective cell migration^43,44^, and, consistent with our findings, P-cadherin depletion in human keratinocytes results in higher cortical tension and reduced differentiation^45^. Additionally, E-cadherin has recently been linked to decreased cortical contractility and low ability of epithelial cells to prevent multipolar divisions, possibly involving Discoidin Domain Receptor 1 (DDR1)-mediated induction of RhoE, which counteracts ROCK^46^. While we found E-Cadherin expression in keratinocytes to be Par3-independent^40^, it is tempting to speculate that elevated P-cadherin instead may serve a similar role. Interestingly, in line with above epithelial carcinoma study^46^, Par3-deficient keratinocytes display both increased expression and junctional localization of DDR1 (Suppl.Fig.2d,f-i), thus correlating with increased P-cadherin levels but inversely correlating with pMLC2 (Suppl.Fig.2f). Though requiring further investigation, these data open the possibility that Par3 mediates actomyosin contraction through restricting P-cadherin and/or DDR1 function, thereby shaping a mechanical force platform at epidermal cell-cell contacts to regulate keratinocyte dynamics.

Although our findings clearly implicate defective mitosis in the increased epidermal differentiation, they do not exclude that additional mechanisms such as local force coupling between non-mitotic and mitotic neighbors^45^ further contribute to premature differentiation upon Par3 loss. Finally, given the plethora of effects that DNA damage can yield in both tissue and cancer stem cells^47^, it will be interesting to dissect in the future if mechanisms similar to those identified here underlie Par3- and/or aPKCλ-dependent skin tumorigenesis^28,48^.

Collectively, this study unravels a novel polarity protein-mediated mechanism that governs tissue homeostasis through coupling mechanical forces with mitotic accuracy and genome integrity, thereby ultimately counteracting premature differentiation. Our findings identify mammalian Par3 as a key integrator of Rho/ROCK-driven keratinocyte mechanics to balance self-renewal and differentiation. The mechanisms revealed here thus provide new molecular and cellular insights into how polarity proteins integrate biomechanical signaling and cell division processes that are fundamental to maintain multilayered tissues.

## METHODS

### Mice

Mice with epidermal *Par3* deletion have been described previously (K14CreneoKI^+/wt^;*Par3*^fl/fl^; *Par3*eKO)^11^. For this study, we crossed *Par3*eKO mice onto a C57BL/6 background. Both male and female mice were included, with the male/female ratio comparable in the control and test groups, and for randomization mice of different genotypes were co-housed. Mice were housed and fed according to federal guidelines. All animal experiments were performed according to institutional guidelines and animal licenses of the State Office of North Rhine-Westphalia, Germany.

### Keratinocytes Isolation and Culture

Primary mouse keratinocytes were isolated from epidermis of newborn control and *Par3*eKO mice. To separate epidermis from dermis, whole skins of P0-P3 mice were incubated in 5mg/ml Dispase II (Sigma-Aldrich) diluted in DMEM/HAM’s F12 medium with low Ca^2+^ (50μM) (Biochrom) supplemented with 10% FCS (chelated), penicillin (100U/ml), streptomycin (100μg/ml, Biochrom #A2212), adenine (1.8×10^-4^M, SIGMA #A3159), l-glutamine (2mM, Biochrom #K0282), hydrocortisone (0.5μg/ml, Sigma #H4001), epidermal growth factor (10ng/ml, Sigma #E9644), cholera enterotoxin (10^-10^M, Sigma #C-8052), insulin (5μg/ml, Sigma #I1882), and ascorbic acid (0.05mg/ml, Sigma #A4034) at 4°C overnight (FAD medium). After incubating the epidermis for 20min in TrypLE (Gibco) at room temperature, dissociated cells were collected and cultured in FAD medium on collagen-I coated plates. Mixed litters were used to isolate control and *Par3*eKO cells.

### Live cell imaging

Briefly, primary keratinocytes isolated from H2B-GFP epidermal *Par3*KO (K14CreneoKI^wt/+^;*Par3*^fl/fl^;H2B-GFP^40^) and control mice (*Par3*^fl/fl^;H2B-GFP) were seeded in μ-Slides (8 Well Glass Bottom ibiTreat; IBIDI) and calcium-switched for 24 hours. In experiments with restoration of contractility, medium containing DMSO (vehicle control) or 1nM Calyculin A (CalA) were added 2 hours before imaging. Time-lapse microscopy was performed using a Leica^®^ DMI 6000 equipped with a Pecon^®^ PM2000 incubator and a PlanApo 20x 0.75 NA objective. Cells were maintained at 37°C, 5% CO2, and time-lapse images were captured every 5 or 10 min for 16 hours. Subsequent analysis of the material was performed with LAS X software (Leica).

### Immunocytochemistry

Cells grown on permanox, glass chamber slides, or Transwell filters were fixed in 4% paraformaldehyde or EtOH/acetone, permeabilized with 0.5% Triton-X (if PFA fixed) in PBS^++^ (0.1mM Mg^2+^ and Ca^2+^) and blocked in 5% bovine serum albumin (BSA). Samples were incubated overnight in primary antibodies diluted in 5% AB buffer (10mM Tris, 150mM NaCl, 0.1%BSA, 0.02% sodium azide), followed by washing and incubation in secondary antibodies for 1h at room temperature. Finally, samples were mounted in Mowiol.

### Immunohistochemistry

For immunofluorescence staining of tissues, paraffin and cryo-sections were used. Paraffin sections were deparaffinized, and antigens were retrieved by boiling in Dako antigen retrieval agent pH9 (Dako) for 20min. Cryo-sections were PFA fixed. After PBS-washing, blocking was performed for 1hr with 5% BSA in PBS at RT. Primary antibodies diluted in either AB buffer or Dako antibody diluent were applied O/N at 4°C in a humidified chamber. AlexaFluor-conjugated secondary antibodies and DAPI (Invitrogen, Darmstadt, Germany) were incubated for 1h at RT before washing and mounting.

### Preparation of cell extracts, SDS-PAGE, and immunoblotting

Analysis of cell extracts was performed as previously described^40^. Briefly, total cell lysates were generated by boiling cells in crude lysis buffer (10mM EDTA, 1% SDS), protein concentrations were determined via BCA assay (Pierce, Thermo Fisher Scientific Darmstadt, Germany), and SDS-PAGE (8-12%PAA) and immunoblotting was performed according to standard procedures.

### Analysis of mitotic duration and aberrant divisions

Time-lapse microscopy videos were used to analyze the mode of division, and divisions were categorized into normal division, lagging strands, tripolar divisions and division with incomplete cytokinesis based on chromatin visualization by GFP-tagged histone H2B and differential interference contrast (DIC). Mitotic duration was analyzed using time-lapse microscopy videos and defined as the difference between the mitotic entry frame and the first frame in cytokinesis. The frame of mitotic entry was defined as the first frame with a nucleus showing early prophase characteristics, such as initiation of chromatin condensation. The frame of cytokinesis was defined as the first frame with visible membrane reestablishment using DIC. Data from three individual experiments was pooled and used to calculate the mean event duration.

### Interphase fluorescence in situ hybridization (iFISH)

Cells were trypsinized, and hypotonic lysis was achieved by resuspension in 75mM KCl for 15 minutes at 37°C, followed by fixation in Carnoy’s fixative (75% methanol: 25% acetic acid) twice for 20 minutes at room temperature before storing samples at −20°C until further use. Genomic DNA and iFISH probes (Tlk2 11qE1/Aurka 2qH3 mouse probe, Leica) were denatured at 73°C for 3 minutes, and hybridization was performed at 37°C overnight. Cells were subsequently washed with 0.4x saline-sodium citrate (SSC) (0.05% Tween-20) for 2 minutes at 68°C and washed with 2xSSC (0.05% Tween-20) at room temperature for 6 minutes. Samples were dehydrated in ethanol series (70-90-100%), air-dried and mounted with VectaShield (Vector Laboratories) containing DAPI.

### Microscopy

Confocal images were acquired with a Spinning Disk microscope (PerkinElmer UltraVIEW, Nikon) using a PlanApo 40x 0.95 NA air objective and Zeiss Meta 710 using a Plan-Apochromat 63x/1.4 NA oil. Epifluorescence images were obtained with a Leica DMI6000 and the following objectives: PlanApo 63x, 1.4 NA; PlanApo 20x, 0.75 NA.

### Particle image velocimetry (PIV)

Particle image velocimetry was performed on 8-bit images of DIC/H2B-GFP from time-lapse micrograph sequences using PIVlab^18^ in MATLAB. Images were pre-processed using the *Contrast-limited adaptive histogram equalization* (*CLAHE*) value of 40px and *high pass filter* of 20px. The size of the interrogation window was of 128×128 pixels followed by 3 steps with subsequent half of the size of the previous. A Gaussian 2×3 fit was used for subpixel accuracy. Image pairs (1^st^-2^nd^, 2^nd^-3^rd^, etc.) were cross-correlated to extract the local displacement in between the two frames. Vector outliers were filtered manually as described previously^18^, and after the removal of outliers, missing vectors were replaced by interpolated data from adjacent time points.

### Analysis of intrinsic mitotic spindle geometry

Keratinocytes were cultivated on chamber slides and switched to 1.8mM Ca^2+^ to induce cell-cell contact formation for 48h. After fixation cells were immunostained for pericentrin and alpha-tubulin, and DNA was visualized using DAPI. Cells in metaphase were selected and the two angles α and β between the metaphase plate and the spindle poles were determined in FIJI^49,50^ using pericentrin and α-tubulin staining. The coherency index c was calculated as follows: c=α/β (with: α≤β≤90°). For spindle shape analysis, thresholds for alpha-tubulin signals were set in FIJI (via *AutoThreshold*) and spindles identified manually with the W*and Tool*. Measurements were performed with the *analyze particles* plugin.

### Quantification of protein signals in immunoblot analyses

The band intensity of non-saturated Western blot signals was determined using the FIJI^49,50^ rectangular tool on digital tiff-files of scanned Western blots. Bar diagrams show the relative protein or phosphorylation signals after normalization using loading controls as indicated.

### Quantification of p-MLC2 junctional intensity

A custom FIJI^49,50^ macro was used to draw constant lines of 15μm length and 50μm width across p-MLC2 junctional sites. The mean grey values were used for quantification.

### Atomic Force Microscopy

Force spectroscopy was performed using a JPK Nanowizard Life Science instrument (JPK) mounted on an inverted optical microscope (Axiovert 200; Zeiss) for sample observation as described previously^51^. Briefly, 5×10^5^ keratinocytes were seeded in 3.5mm diameter plates (TPP) pre-coated with collagen-I, and switched to 1.8mM Ca^2+^ to induce cell-cell contact formation for 48h. For nano-indentation experiments, silicon nitride pyramidal cantilevers (MLCT, Bruker Daltonics) with a nominal spring constant of 0.03 Nm^-1^, or spherical silicon dioxide beads with a diameter of 3.5 μm glued onto tip-less silicon nitride cantilevers (NanoAndMore/ sQube, CP-PNPL-SiO-B-5) with a nominal spring constant of 0.08 Nm^-1^ were used. The cantilever speed was kept constant at *v*=1.5μm/s, and the force set-point was 0.05nN. Data analysis was performed with Atomic J^52^ and data fitted as described previously^51^.

### UV irradiation

Confluent primary keratinocyte cultures were washed once with PBS, then covered with PBS and irradiated with a dose of 100mJ/cm^2^ UV-B (304nm). After irradiation, PBS was removed and cells were incubated at standard culture conditions for the indicated time points before protein lysates were collected.

### Epithelial sheet contraction assay

5×10^5^ keratinocytes were plated in triplicates on 6-well culture plates. 24 hours after seeding, confluent cultures were switched to high calcium for 48h. Cells were subsequently washed with PBS^++^ and incubated with 2ml Dispase II (2.4U/ml in FAD medium with 1.8mM CaCl_2_) (Roche) per well, for 30 min at 37°C, to detach the keratinocyte sheets. Remaining contacts at the sides of the wells were removed using a scalpel. Pharmacologic compounds were pre-incubated for 24 hours prior to the assay and were present during dispase-mediated lift. Images of resulting contracted keratinocyte sheets were acquired using a GelDOC XR+ (BioRad) station, and analyzed in FIJI^49,50^.

### Stratified primary murine keratinocyte cultures

0.8-1.2×10^5^ primary keratinocytes were seeded onto porous, collagen-I-coated TransWell filters (Corning, #3470, 0.4μm pore size). 24 hours after seeding confluent cultures were switched to high calcium and subsequent stratification was allowed during a 10 days-culture period. In case of inhibitor treatment, compounds were added at the time of calcium switch and replaced every second day.

### Slot blot

Primary keratinocytes were cultured and UV-B irradiated using 3mJ/cm^2^ UV-B as described before. Cells were lysed using lysis buffer (0.2% SDS, 200mM NaCl, 5mM EDTA, 100mM Tris-HCl, pH8.5) and genomic DNA was extracted by isopropanol extraction. DNA was denatured for 5min at 95°C, put directly on ice, and blotted onto Amersham Hybibd-N+ membrane (RPN119B GE Healthcare) using a Whatman 96-well slot blotting device at 300 mbar vacuum. Cross-linking of the DNA was carried out for 2 h at 80 °C. The membrane was blocked for 30min in 3% milk-PBS-T, incubated with anti-CPD antibodies (TDM-2, Cosmo Bio 1:10.000) overnight at 4°C, and peroxidase-conjugated secondary antibodies (Jackson Immuno Research 1:10.000) for 1h. After incubation with ECL (RPN2232, GE Healthcare), DNA Lesions were visualized and quantified on the Molecular Imager Gel Doc Imaging System (Bio-Rad) using Image Lab 5.0.

### Antibodies

Detailed information on the antibodies used in this study is provided in Supplementary Table 1.

### Statistical analyses

Statistical analyses were performed using GraphPad Prism software (GraphPad, version 6.0). Significance was determined by Mann–Whitney U-test, Student’s *t*-test, Kruskal–Wallis ANOVA with Dunn’s post-hoc test, One-Way ANOVA and Two-Way ANOVA with Tuckey multiple comparison or Sidak’s multiple comparisons test respectively as indicated in the figure legends. All data sets were subjected to normality tests by D’Agostino-Pearson omnibus test. For statistical analysis of mitotic duration a mixed effects model was used in RStudio. P-values are ranged as follows: *, p<0.05; **, p<0.01; ***, p<0.001; as detailed in the figure legends.

## ACKNOWLEDGEMENTS

We thank Katherine Dodel for technical assistance, and the CECAD imaging facility and the UoC animal facilities for important services. We thank the Iden, Niessen, Wickström, and Bazzi laboratories as well as Christian Reinhardt and Jan Hoeijmakers for helpful discussions. Furthermore, we are grateful to Floris Foijer for advice on iFISH, and to Mirka Uhlirova for various discussions and critical reading of the manuscript. We acknowledge Walter Birchmeier and Shigeo Ohno for sharing mouse lines. This project was funded by the German Research Foundation (DFG) (grants SPP1782-ID79/2-1, and CRC829/A10). Work in the Iden laboratory is further supported by the Excellence Initiative of the German federal and state governments (CECAD) and Center for Molecular Medicine Cologne (CMMC).

## AUTHOR CONTRIBUTIONS

Conceptualization, methodology, and validation: M.D.G., S.L., S.I.; investigation: S.L., M.D.G., M.S., S.I.; formal analysis: M.D.G., S.L., M.S., S.I.; resources: S.I.; visualization: M.D.G., S.L.; writing (original draft): S.L., M.D.G.; writing (review and editing): S.I.; supervision and funding acquisition: S.I..

